# Ribosome protein mutant cells rely on the GR64 cluster of gustatory receptors for survival and proteostasis in *Drosophila*

**DOI:** 10.1101/2021.05.10.443504

**Authors:** Michael E. Baumgartner, Iwo Kucinski, Eugenia Piddini

## Abstract

Mutations in ribosome protein (*Rp*) genes and ribosome biogenesis factors result in debilitating diseases known as ribosomopathies. Recent studies in *Drosophila* have shown that cells heterozygous mutant for *Rp* genes (*Rp/+*) exhibit proteotoxic stress and aggregates, which drive stress pathway activation and apoptosis. Understanding how *Rp/+* cells fend off proteotoxic stress could suggest mechanisms to ameliorate these and other conditions caused by proteotoxic stress. Here we find that *Rp/+* epithelial cells express all six Gustatory Receptor 64 (Gr64) genes, a cluster of sugar receptors involved in taste sensation. We show that *Rp/+* cells depend on Gr64 for survival and that loss of Gr64 autonomously exacerbates stress pathway activation and proteotoxic stress by negatively effecting autophagy and proteasome function in *Rp/+* cells. This work identifies a non-canonical role in proteostasis maintenance for a family of gustatory receptors known for their function in neuronal sensation.

## Introduction

Mutations in ribosomal proteins or ribosome biogenesis factors can result in a class of disorders known as ribosomopathies. While different mutations often yield different clinical manifestations, individuals with ribosomopathies typically present with deficiencies in haematopoiesis, defects in neural crest-derived tissues, and an increased risk of cancer (Armistead & Triggs-Raine, 2014; Aspesi & Ellis, 2019; Mills & Green, 2017). While roles have been established for nucleolar stress, p53 activation, and translational defects in disease progression, ribosomopathy etiology remains poorly understood (Armistead & Triggs-Raine, 2014).

Ribosome protein mutations are well studied in *Drosophila*, wherein the majority of cytosolic ribosome protein genes yield the so-called ‘Minute’ phenotype when heterozygous mutant (Marygold et al., 2007). *Rp/+* flies are viable and fertile but exhibit a developmental delay (Bridges & Morgan, 1923). Epithelia in *Rp/+* larvae exhibit reduced translation rates, increased cell-autonomous apoptosis, and stress pathway activation, including the p53/DNA damage response, JNK, JAK/STAT, Toll/IMD signalling, Xrp1/Irbp18, and the oxidative stress response (Baillon et al., 2018; Blanco et al., 2020; Kucinski et al., 2017; Lee et al., 2018; Meyer et al., 2014). *Rp/+* cells also undergo cell competition when confronted with wildtype cells in mosaic tissues: *Rp/+* cells exhibit a growth defect and, when proximal to wildtype cells, undergo apoptosis and are eliminated from the tissue (Martín et al., 2009; Morata & Ripoll, 1975). *Rp/+* cells are therefore said to behave as ‘losers’ relative to wildtype ‘winners’.

*Rp/+* cells have recently been shown to suffer from chronic proteotoxic stress. They exhibit proteasome and autophagy defects, activation of the integrated stress response (ISR), a stoichiometric imbalance of large and small ribosomal subunit proteins, and an accumulation of intracellular protein aggregates (Albert et al., 2019; Baumgartner et al., 2021; Recasens-Alvarez et al., 2021; Tye et al., 2019). Importantly, boosting proteostasis rescues stress pathway activation, cell autonomous apoptosis, and competitive elimination(Baumgartner et al., 2021; Recasens-Alvarez et al., 2021). These findings point to proteotoxic stress as a potential driver of the pathologies associated with ribosomopathies.

Gustatory Receptors 64 are a group of six tandem gustatory receptor genes (a through f) involved in mediating sensation of sugars, fatty acids, and glycerol in the adult nervous system (Fujii et al., 2015; Kim et al., 2018; Miyamoto et al., 2013; Slone et al., 2007). *Gr64*’s are thought to sense distinct ligands via distinct mechanisms: *Gr64e*, for instance, is reported to act as a ligand gated ion channel in response to glycerol binding, whereas it acts downstream of Phospholipase C in fatty acid sensation (Kim et al., 2018, p. 64). Gr64’s are also thought to function as heterodimers with each other and alternate gustatory receptors, as, for instance, both Gr64a and Gr64f are required to mediate responses to certain sugars (Jiao et al., 2008).

By characterising the biology of *Rp/+* cells, we identify a new function for *Gr64* genes in proteostasis maintenance in epithelial cells. We find that loss of Gr64 drives substantial apoptosis in non-competing and competing *Rp/+* cells and exacerbates stress pathway activation. Loss of Gr64 furthermore exacerbates proteotoxic stress in *Rp/+* cells, likely by reducing proteasome and autophagy function. *Rp/+* cells are therefore acutely reliant on *Gr64* in order to manage proteotoxic stress and, thus, for their survival.

## Results and Discussion

Mining the list of genes differentially expressed in cells heterozygous mutant for a ribosomal protein, *RpS3*, we observed an upregulation of all six Gr64 gustatory receptors relative to wildtype (Figure 1a and Kucinski et al., 2017). Gr64s were also upregulated in cells mutant in *mahj*^*-/-*^, an E3 ubiquitin ligase, whose mutation also leads to proteotoxic stress and cell competition (Baumgartner et al., 2021; Kucinski et al., 2017; Tamori et al., 2010). This was conspicuous, as *Gr64* has no known non-neuronal nor larval function. In order to explore the role of *Gr64s* in *Rp/+* cells, we tested the effect of removing one copy of the *Gr64* locus on *RpS3*^*+/-*^ larvae, using a deficiency spanning the Gr64 locus, along with rescuing constructs of other affected genes (*ΔGr64*) (Slone et al., 2007). *ΔGr64/ΔGr64* flies present with no known phenotypes other than a deficient gustatory responses (Slone et al., 2007). *RpS3*^*+/-*^, *ΔGr64*^*+/-*^ wing discs, however, exhibited a marked increase in apoptosis over levels seen in wing discs carrying either mutation alone (Figure 1b-d, Figure 1-figure supplement 1), indicating that *RpS3*^*+/-*^ cells are acutely reliant on *Gr64* for their survival. We confirmed this result using a precise CRISPR/Cas9 deletion of the Gr64 cluster, *Gr64af* (Kim et al., 2018) (Figure 1-figure supplement 2). *Gr64* does not contribute to survival in the non-Minute context, as non-Minute wing discs homozygous null for *Gr64* exhibited no increase in apoptosis relative to the wildtype (Figure1-figure supplement 3). To determine if this reflects a systemic or cell autonomous role for *Gr64*, we knocked down Gr64 specifically in the posterior compartment, using the *hedgehog-Gal4* driver. We used an RNAi line against Gr64f, which, given that the *Gr64* cluster locus is polycistronic (Slone et al., 2007), is likely to silence multiple Gr64s. Gr64f-RNAi expression in wildtype discs yielded no appreciable change in levels of apoptosis (Figure 1e), whereas expression of Gr64f-RNAi in *RpS3*^*+/-*^ wing discs yielded a strong increase in apoptosis (Figure 1f-g), indicating that *RpS3*^*+/-*^ cells are cell autonomously dependent on Gr64 for their survival.

**Figure 1:**
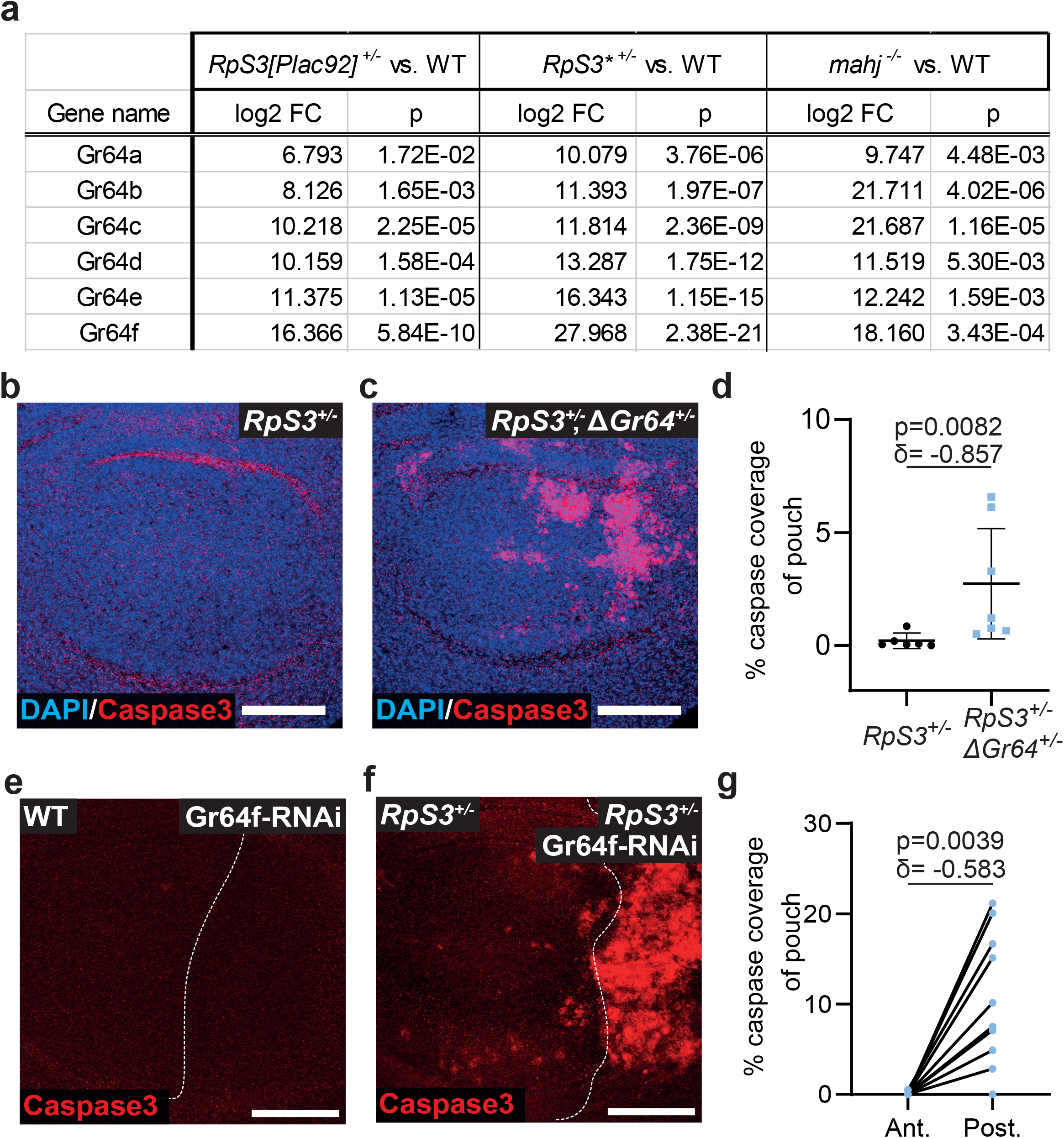
Non-competing *RpS3*^*+/-*^ cells depend on Gr64 for their survival. (**a**) Log 2 fold change in Gr64 transcript expression relative to wildtype in wing discs heterozygous mutant for *RpS3* (as measured from two separate mutant alleles: *RpS3[Plac92]* or *RpS3\**) or homozygous mutant for *Mahjong*. Numbers and p-values are derived from Kucinski et al., 2017. (**b**-**d**) Wing discs heterozygous mutant for *RpS3* without (**b**) or with (**c**) a heterozygous deficiency in the *Gr64* locus (ΔGr64) and assessed for cell death with a staining for cleaved-caspase3 (red), along with quantification in (**d**) (n_*RpS3*_=6, n_*RpS3*,ΔGr64_=7, two-sided Mann-Whitney U test with Cliff’s δ effect size metric, the horizontal line indicates the mean and the whiskers reflect the 95% confidence interval.). (**e**-**g**) Wildtype (**e**) or *RpS3*^*+/-*^ (**f**) wing discs expressing Gr64f-RNAi in the posterior compartment with hh-Gal4 and assessed for cell death with a staining for cleaved caspase3 (red) along with quantification in (**g**) (n=12, two-sided Wilcoxon signed rank test with Cliff’s δ effect size metric). For this and all other figures, scale bars correspond to 50µm, white dashed lines denote the compartment boundaries, the anterior compartment is shown on the left side of the image and dorsal is up.

Having established a pro-survival role for *Gr64* in non-competing *RpS3*^*+/-*^ cells, we then tested whether *Gr64*’s contribute to the survival of competing *RpS3*^*+/-*^ losers. We therefore generated *RpS3*^*+/-*^ losers competing against *RpS3*^*+/+*^ winners in wing discs carrying heterozygous mutations in one of any of the six Gr64 genes(Yavuz et al., 2015) (Figure 2). Strikingly, heterozygousity for any *Gr64* yielded a substantial increase in *RpS3*^*+/-*^ loser cell death at the winner/loser interface (Figure 2h). Furthermore, loser cell clones were smaller in wing discs carrying mutations in *Gr64b, Gr64c, Gr64d, Gr64e*, or *Gr64f* (Figure 2i). *RpS3*^*+/-*^ loser cells are therefore dependent on *Gr64*s in both non-competitive and competitive conditions.

**Figure 2:**
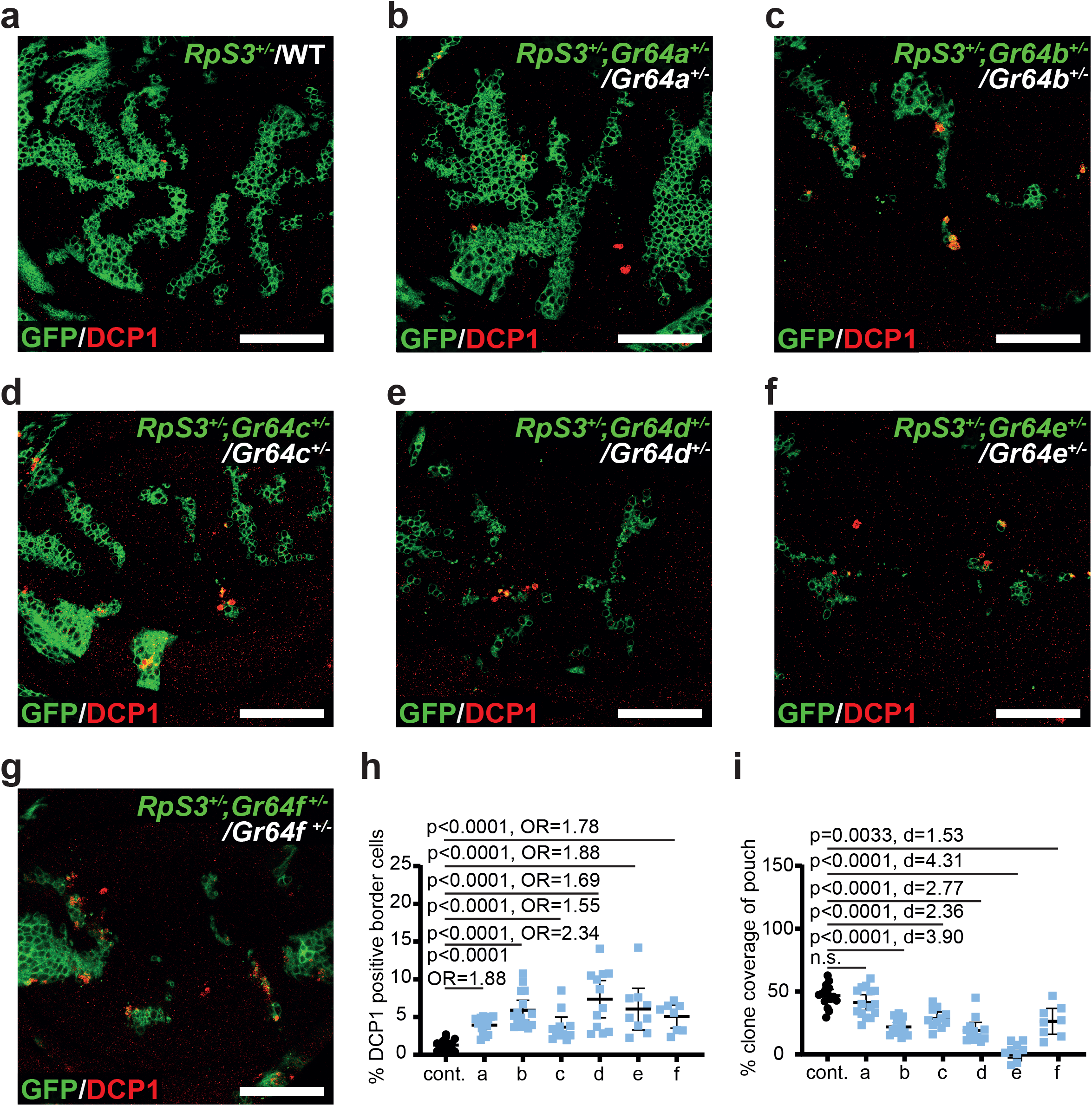
Heterozygosity at Gr64 loci exacerbates competitive *RpS3*^*+/-*^ loser cell elimination. (**a**-**g**) Representative images of wing discs containing *RpS3*^*+/-*^ losers (green) competing against wildtype winners (unlabelled) and stained for cleaved-DCP1 (red). *RpS3*^*+/-*^ clones were generated in a wildtype background (**a**) or in wing discs heterozygous for any one of the Gr64 genes a through f (**b**-**g**). (**h**) Quantification of the percentage of cells undergoing apoptosis at the loser clone border in the wing discs shown in (**a**-**g**). Statistics reflect multiple logistic regression across 3 replicates (details provided in methods) (**i**) Quantification of loser cell growth in the wing discs shown in (**a**-**g**), as measured by the percent loser clone coverage of the pouch. Statistics reflect student’s t-test with FDR p-correction and cohen’s d effect size metric. n_control_=16, n_Gr64a_=15, n_Gr64b_=15, n_Gr64c_=11, n_Gr64d_=12, n_Gr64e_=9, n_Gr64f_=8. For all quantifications, the horizontal line indicates the mean and the whiskers reflect the 95% confidence interval.

*Rp/+* mutations cause stress pathway activation and cellular malfunctions, many of which are linked with the loser status (Albert et al., 2019; Baumgartner et al., 2021; Ellis, 2014; Kucinski et al., 2017; Lee et al., 2018; Recasens-Alvarez et al., 2021; Tye et al., 2019). To evaluate how *Gr64* loss affects *Rp/+* stress responses, we identified milder expression conditions for Gr64f RNAi, using the posterior compartment-specific *engrailed-Gal4* driver. In these conditions Gr64f knockdown in non-competing *RpS3*^*+/-*^ cells exhibited a comparatively mild increase in apoptosis (Figure 3a-b), allowing us to investigate *Gr64* function without widespread cell death. Gr64f-RNAi yielded increased activation of the *Nrf2*/oxidative stress response, as measured by *GstD1-GFP* reporter expression(Sykiotis & Bohmann, 2008) (Figure 3c-d). Gr64f-RNAi did not yield an increase in *GstD1-GFP* signal in wildtype wing discs (Figure 3-figure supplement 1a-b), demonstrating that this result is specific to the *RpS3*^*+/-*^ context. Consistent with these results, an increase in *GstD1-GFP* expression was also observed in *RpS3*^*+/-*^ wing discs heterozygous for *ΔGr64* (Figure 3-figure supplement 1c-e). Furthermore, Gr64f-RNAi in *RpS3*^*+/-*^ resulted in a mild increase in JNK pathway and ISR activity, as measured by immunostaining for phosphorylated JNK (Figure 3e-f) and phosphorylated eIF2α (Figure 3g-h), respectively. No differences in phospho-JNK (Figure 3-figure supplement 2 a-b) or phospho-eIF2α signal (Figure 3-figure supplement 2c-d) levels were seen in the posterior compartments of otherwise genetically identical wing discs lacking the Gr64f RNAi, confirming that this result is due to RNAi expression. These data indicate that loss of Gr64 exacerbates stress responses seen in *Rp/+* cells.

**Figure 3:**
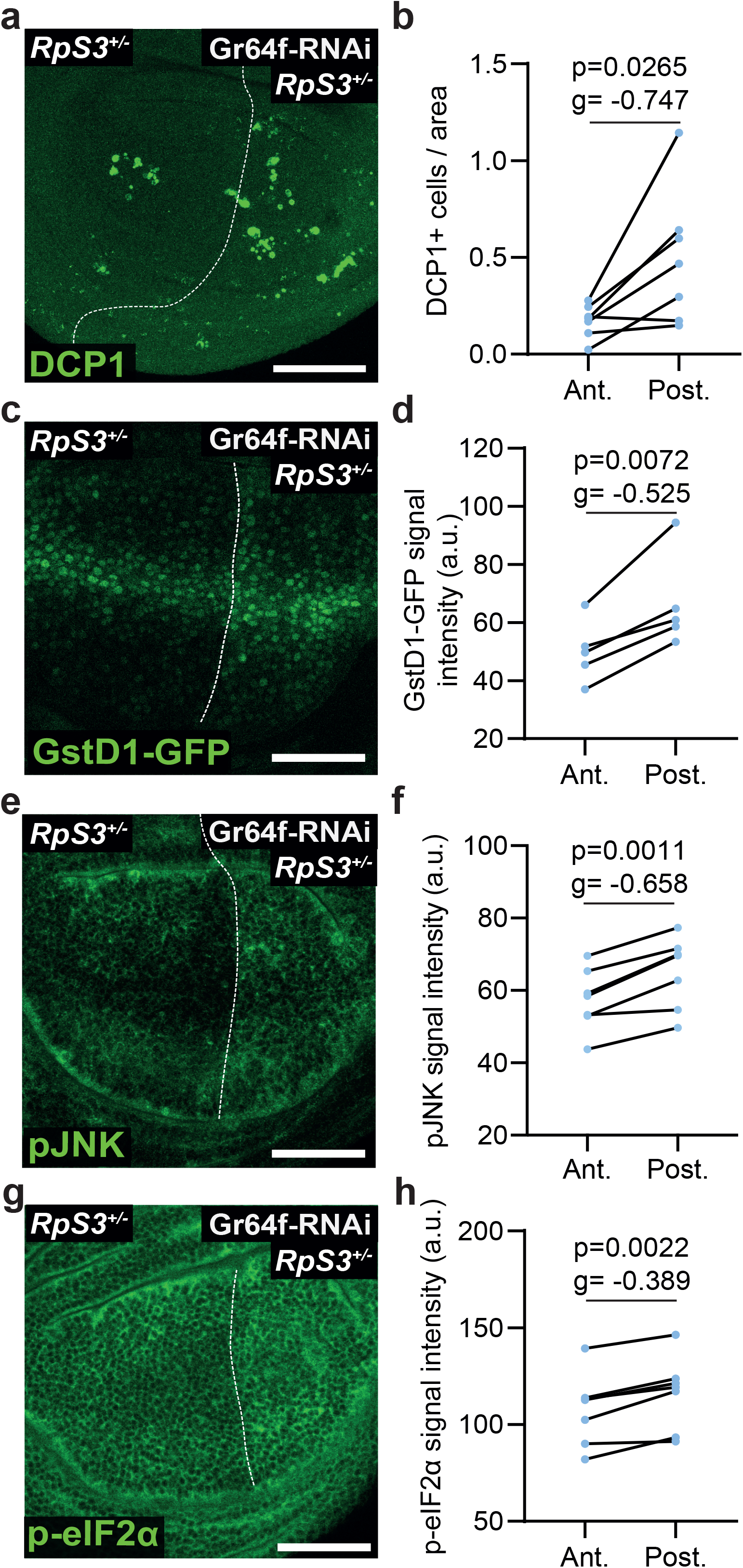
Loss of Gr64 worsens loser-associated stress pathway activation in *RpS3*^*+/-*^. (**a**-**b**) *RpS3*^*+/-*^ wing discs expressing Gr64f-RNAi in the posterior compartment stained for cell death as measured by cleaved DCP1 (green) (**a**) along with quantification in (**b**) (n=7, two-sided Wilcoxon signed rank test with Cliff’s δ effect size metric). (**c**-**d**) *RpS3*^*+/-*^ wing discs expressing Gr64f-RNAi in the posterior compartment and assessed using the GstD1-GFP reporter (green) (**c**) along with quantification in (**d**) (n=6, two-sided paired t-test with Hedges’ g effect size metric). (**e**-**f**) *RpS3*^*+/-*^ wing discs expressing Gr64f-RNAi in the posterior compartment and stained for phosphorylated JNK (green) (**e**) along with quantification in (**f**) (n=7, two-sided paired t-test with Hedges’ g effect size metric). (**g-h**) *RpS3*^*+/-*^ wing discs expressing Gr64f-RNAi in the posterior compartment and stained for phosphorylated eIF2α (green) (**g**) along with quantification in (**h**) (n=7, two-sided paired t-test with Hedges’ g effect size metric).

We next investigated how *Gr64* alleviates stress signalling and improves viability of *Rp/+* cells. We previously reported that ISR and oxidative stress response in *RpS3*^*+/-*^ cells are driven by accumulation of protein aggregates, accompanied by reduced proteasome and autophagy flux, leading to sustained proteotoxic stress (Baumgartner et al., 2021). Thus, we tested the effect of Gr64 reduction on proteostasis pathways. Using ProteoFlux and ReFlux, genetic pulse-chase reporters of proteasome and autophagy flux respectively (Baumgartner et al., 2021), we found that Gr64f-RNAi results in a further reduction in flux through the proteasome (Figure 4a-c) and autophagosome (Figure 4d-f), beyond the defects already present in *RpS3*^*+/-*^. These data indicate that loss of Gr64 yields a substantial impairment in cellular proteolytic pathways. Furthermore, Gr64f-RNAi yields an increase in ref(2)P-positive foci (Figure 4g), structures which we have previously shown to co-localize with poly-ubiquitinylated aggregates in *RpS3*^*+/-*^ cells (Baumgartner et al., 2021). We conclude that Gr64 loss causes increased stress signalling and cell death by exacerbating proteotoxic stress. Worsening of proteotoxic stress in ribosome mutants could derive from an inhibition of protein catabolic processes or from an increase in protein translation, which by producing more proteins could cause a further burden on proteostasis. Interestingly, OPP, a global translation reporter, revealed a mild but statistically significant decrease in translation upon expression of Gr64f-RNAi (Figure 4h-i). This is consistent with the observed increase in eIF2α-phosphorylation (Figure 3g-h), which is a translational inhibitor as well as a stress pathway marker (Pakos-Zebrucka et al., 2016). Thus, the increase in proteotoxic stress observed upon Gr64 reduction is not due to increased translation but rather to a failure to clear defective proteins.

**Figure 4:**
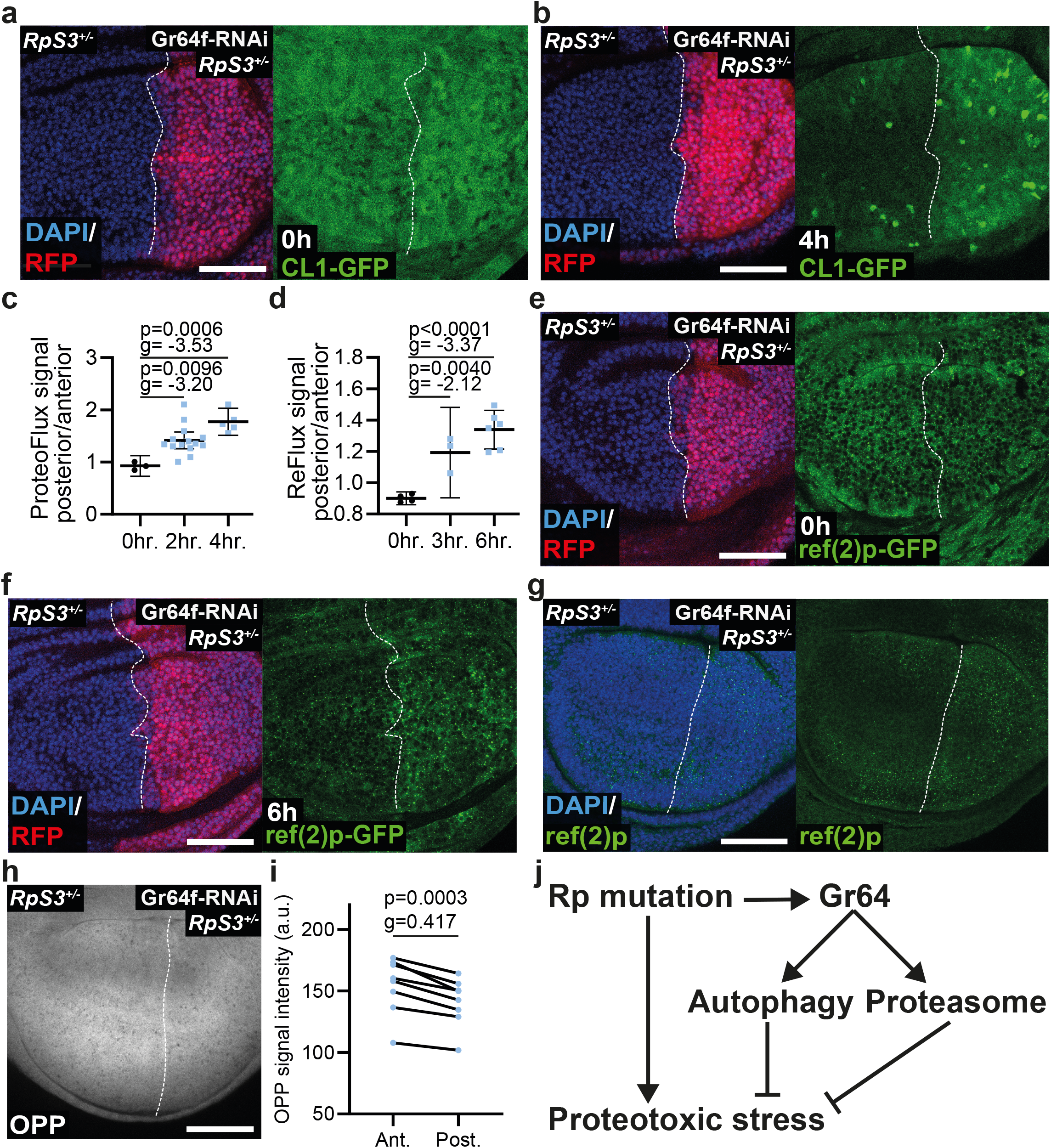
Loss of Gr64 exacerbates proteostasis defects in *RpS3*^*+/-*^ by impairing cellular catabolism. (**a**-**c**) *RpS3*^*+/-*^ wing discs expressing Gr64f-RNAi in the posterior compartment, marked with RFP (red) and expressing the CL1-GFP/ProteoFlux construct (green) 0 (**a**) or 4 (**b**) hours after heat shock along with quantification in (**c**) (n_0hr_=3, n_2hr_=14, n_4hr_=5, two-sided paired t-test with Hedges’ g effect size, the horizontal line indicates the mean and the whiskers reflect the 95% confidence interval). (**d**-**f**) *RpS3*^*+/-*^ wing discs expressing Gr64f-RNAi in the posterior compartment, marked with RFP (red) and expressing the ref(2)P-GFP/ReFlux construct (green) 0 (**e**) or 6 (**f**) hours after heat shock along with quantification in (**d**) (n_0hr_=4, n_3hr_=3, n_6hr_=6, two-sided paired t-test with Hedges’ g effect size, the horizontal line indicates the mean and the whiskers reflect the 95% confidence interval). (**g**) *RpS3*^*+/-*^ wing discs expressing Gr64f-RNAi in the posterior compartment stained for ref(2)P (green). (**h**-**i**) *RpS3*^*+/-*^ wing discs expressing Gr64f-RNAi in the posterior compartment and assessed for translation via OPP (grey) (**h**) along with quantification in (**i**) (n=8, two-sided paired t-test with Hedges’ g effect size metric). (**j**) Proposed model: heterozygous mutation in a ribosomal protein mutant gene triggers proteotoxic stress, potentially via a stoichiometric imbalance of ribosomal proteins. The *Rp/+* cell is then highly dependent upon the autophagosome and proteasome to perform proteolytic functions. Gr64 contributes to survival and alleviates stress pathway activity by promoting proteolysis.

In this study, we have identified Gr64 taste receptors as novel players in proteostasis control and as cytoprotective regulators in epithelial cells affected by proteotoxic stress. How Gr64s contribute to proteostasis is unclear. Our data rule out a role of Gr64 in repressing translation and are instead consistent with a function in modulating protein catabolism (proposed model in Figure 4j). This is potentially mediated by calcium signalling, as Gr64 activity typically induces calcium release and calcium is involved in protein folding, the integrated stress response, proteasome and autophagy function (Bootman et al., 2018; Carreras-Sureda et al., 2018; Decuypere et al., 2011; Mukherjee et al., 2017).

Taste receptors have been implicated in non-taste-related chemo-sensation in neuronal and neuroendocrine cells both in flies and in mammals (Fujii et al., 2015; Miyamoto & Amrein, 2014; Park & Kwon, 2011; Shirazi-Beechey et al., 2014). However, a role in proteostasis and a function in epithelia have not previously been described for taste receptors. Interestingly, dysregulation of olfactory and gustatory receptors has been observed in non-olfactory human brain tissue from individuals suffering from protein-aggregate driven neurodegenerative disorders, including Alzheimer’s disease, Parkinson’s disease, and Creutzfeld-Jacob disease as well as in a mouse model of Alzheimer’s (Ferrer et al., 2016), and olfactory receptors are expressed near to amyloid plaques in a mouse model of Alzheimer’s (Gaudel et al., 2019). It is therefore possible that gustatory and olfactory receptors play a conserved role in proteostasis.

## Acknowledgements

We would like to thank the Amrein lab and S.J. Moon for providing us with their many *Gr64 Drosophila* lines. We would also like to thank T.E. Rusten for graciously providing us with the ref(2)P antibody. We would like to thank the Wolfson Bioimaging Facility at the University of Bristol for access to their microscopes. This work was supported a Cancer Research UK Programme Foundation Award to E.P. (Grant C38607/A26831) and a Wellcome Trust Senior Research Fellowship to E.P. (205010/Z/, 16/Z).

## Materials and Methods

### Fly husbandry and stocks

*Drosophila* lines were kept in an incubator set to 25°C and reared on food prepared according to the following recipe: 7.5 g/L agar powder, 50 g/L baker’s yeast, 55 g/L glucose, 35 g/L wheat flour, 2.5% nipagin, 0.4% propionic acid and 1.0% penicillin/streptomycin. All larvae were dissected at the wandering third instar stage. For experiments with heat-shock-induced clones, vials were transferred to a water bath set to 37°C for 25 minutes on day three after egg laying. The vials were then immediately returned to the 25°C incubator and allowed to grow as normal for three more days prior to dissection. Only female larvae were dissected.

The following *Drosophila melanogaster* lines were obtained from the Bloomington Drosophila Resource Center: *RpS3[Plac92]* (Cat#BL5627), *en-Gal4, UAS-RFP* (Cat#BL30557), *en-Gal4* (Cat#BL30564). The Gr64f-RNAi line (Cat#v100156) was obtained from the Vienna Drosophila Resource Centre. The *hs-CL1-GFP (ProteoFlux), hs-ref(2)P-GFP(ReFlux)* and *hs-FLP, UAS-CD8-GFP;; FRT82B, RpS3[Plac92], act>RpS3>Gal4/TM6b* were reported in (Baumgartner et al., 2021). The following lines were kindly provided by Hubert Amrein: R1; R2; ΔGr64 (R1 and R2 are constructs rescuing all other genes flanking the Gr64 locus affected by the deletion) (Slone et al., 2007), Gr64a-Gal4, Gr64b-lexA, Gr64c-lexA, Gr64d[1], Gr64e-lexA, and Gr64f-lexA. Gr64-Gal4/lexA insertions are both expression reporters and validated null mutants (Fujii et al., 2015; Yavuz et al., 2015). The *hs-FLP;; FRT82B* and *y[1], w[1118]* lines were provided by Daniel St. Johnston. The *hh-Gal4/TM6B* was provided by Jean-Paul Vincent. The *GstD1-GFP* line was described in (Sykiotis & Bohmann, 2008). The *Gr64af* deletion was described in (Kim et al., 2018).

Genotypes for all experimental crosses are provided in the following table:

| Figure/Panel | Genotype |
| --- | --- |
| **Main Figures** |  |
| Figure 1b | *FRT82B*, *RpS3[Plac92]*, *ubi-GFP/+* |
| Figure 1c | *R1/+; R2/+; FRT82B, RpS3[Plac92], ubi-GFP/ΔGr64* |
| Figure 1e | *UAS-Gr64f-RNAi (KK)/GstD1-GFP; hh-Gal4/+* |
| Figure 1f | *UAS-Gr64f-RNAi (KK)/GstD1-GFP; FRT82B, RpS3[Plac92], hh-Gal4/+* |
| Figure 2a | *hs-FLP, UAS-CD8-GFP/+;; RpS3[Plac92], act>RpS3>Gal4/+* |
| Figure 2b | *hs-FLP, UAS-CD8-GFP/+;; RpS3[Plac92], act>RpS3>Gal4/Gr64a-Gal4* |
| Figure 2c | *hs-FLP, UAS-CD8-GFP/+;; RpS3[Plac92], act>RpS3>Gal4/Gr64b-LexA* |
| Figure 2d | *hs-FLP, UAS-CD8-GFP/+;; RpS3[Plac92], act>RpS3>Gal4/Gr64c-LexA* |
| Figure 2e | *hs-FLP, UAS-CD8-GFP/+;; RpS3[Plac92], act>RpS3>Gal4/Gr64d[1]* |
| Figure 2f | *hs-FLP, UAS-CD8-GFP/+;; RpS3[Plac92], act>RpS3>Gal4/Gr64e-LexA* |
| Figure 2g | *hs-FLP, UAS-CD8-GFP/+;; RpS3[Plac92], act>RpS3>Gal4/Gr64f-LexA* |
| Figure 3a | *hs-FLP; en-Gal4/Gr64f-RNAi (KK); FRT82B, RpS3[Plac92], tub>dsRed/Frt82B* |
| Figure 3c | *hs-FLP; en-Gal4, GstD1-GFP/Gr64f-RNAi (KK); FRT82B, RpS3[Plac92], tub>dsRed/Frt82B* |
| Figure 3e | *hs-FLP; en-Gal4/Gr64f-RNAi (KK);FRT82B,* |
|  | *RpS3[Plac92], tub>dsRed/Frt82B* |
| Figure 3g | *hs-FLP; en-Gal4/Gr64f-RNAi (KK);FRT82B, RpS3[Plac92], tub>dsRed/Frt82B* |
| Figure 4a-b | *hs-CL1-GFP (ProteoFlux)/(+ or y); enGal4, UAS-RFP/Gr64f RNAi (KK); FRT82B, RpS3[Plac92], tub>dsRed/+* |
| Figure 4e-f | *hs-ref(2)P-GFP (ReFlux)/(+ or y); enGal4, UAS-RFP/Gr64f RNAi (KK); FRT82B, RpS3[Plac92], tub>dsRed/+* |
| Figure 4g-h | *hs-FLP; en-Gal4/Gr64f-RNAi (KK);FRT82B, RpS3[Plac92], tub>dsRed/Frt82B* |
| **Figure Supplements** |  |
| Figure 1 – figure supplement 1a | *R1/+; R2/+; FRT82B, ubi-GFP/ΔGr64* |
| Figure 1 – figure supplement 1b | *R1/+; R2/+; FRT82B, RpS3[Plac92], ubi-GFP/ΔGr64* |
| Figure 1- figure supplement 2a | *Gr64af/+* |
| Figure 1- figure supplement 2b | *Frt82B, RpS3{Plac92}, tub>dsRed/+* |
| Figure 1- figure supplement 2c | *Gr64af/ Frt82B, RpS3{Plac92}, tub>dsRed* |
| Figure 1- figure supplement 3a | Wildtype |
| Figure 1- figure supplement 3b | *Gr64af/Gr64af* |
| Figure 3 – figure supplement 1a | *UAS-Gr64f-RNAi (KK)/GstD1-GFP; hh-Gal4/+* |
| Figure 3 – figure supplement 1c | *GstD1-GFP/+; FRT82B, RpS3[Plac92], hh-Gal4/+* |
| Figure 3 – figure supplement 1d | *R1/+; R2/GstD1-GFP; FRT82B, RpS3[Plac92],* |
|  | *hh-Gal4/ΔGr64* |
| Figure 3 – figure supplement 2a | *hs-FLP; en-Gal4/+; FRT82B, RpS3[Plac92], tub>dsRed/Frt82B* |
| Figure 3 – figure supplement 2c | *hs-FLP; en-Gal4/+; FRT82B, RpS3[Plac92], tub>dsRed/Frt82B* |

### Immunofluorescence

Larvae at the wandering third instar stage were washed once and then dissected in PBS before being immediately transferred to a pre-chilled vial of PBS. Samples were then fixed in 4% formaldehyde in PBS at room temperature for 20 minutes. Samples were then washed three times in PBS and then permeabilized in 0.25% Triton X-100 in PBS (PBST). The PBST was then aspirated and replaced with blocking buffer (4% foetal calf serum in PBST) and incubated for thirty minutes at room temperature. Primary antibodies were diluted in blocking buffer, and primary incubations took place overnight at 4°C on a rocker. Samples were then washed three times in PBST at room temperature for three minutes, followed by a one-hour incubation with secondary antibody diluted 1:500 in blocking buffer along with 0.5µg/mL 4’,6-diamidino-2-phenylindole (DAPI). Secondary antibodies used were Alexa-Fluor 488, 555, or 633 (Molecular Probes). Samples were then again washed three times for ten minutes in PBST before being mounted in VECTASHIELD (Vector Laboratories) on a borosilicate glass slide (number 1.5, VWR International).

The antibodies used were: Rabbit anti-pJNK pTPpY (1:500, Promega, Cat#V93B), Rat anti-Ci(1:1,000, DSHB, Cat#2A1), Rabbit anti-Ref(2)P (1:2,000, provided by Tor Erik Rusten(Katheder et al., 2017)), Rabbit anti-cleaved Caspase-3 (1:25,000, Abcam, Cat#13847), Rabbit anti-DCP1 (1:2,500, Cell Signalling, Cat#9578S), Rabbit anti-p-eIF2α (1:500, Cell Signalling, Cat#3398T).

### ProteoFlux and ReFlux pulse-chase assays

On day six after egg laying, vials containing third instar larvae carrying the ProteoFlux or ReFlux constructs were transferred to a water bath set to 37°C for 40 or 45 minutes, respectively. Larvae were then immediately dissected and transferred to ice cold 4% formaldehyde in PBS to act as a zero timepoint. The vials were then returned to the 25°C incubator and dissected at 2 and 4 hours after heat shock for ProteoFlux and 3 and 6 hours after heat shock for ReFlux. Larvae were then fixed and mounted as normal.

### OPP translation assay

Larvae were washed once and then dissected in pre-warmed Schneider’s medium. Hemi-larvae were then transferred to a 1.5 mL Eppendorf tube containing 5µM OPP (Molecular Probes, Cat#C10456) diluted in Schneider’s medium and placed in a heating block set to 25°C for 15 minutes. Samples were then washed quickly in PBS and then fixed in 4% formaldehyde in PBS for 20 minutes at room temperature, permeabilized for 30 minutes at room temperature in 0.5% PBST and incubated for 30 minutes in blocking buffer. Samples were then washed in PBS and staining was performed using the Click-iT Plus protocol according to manufacturer’s instructions.

### Imaging, quantification, and statistical analysis

All images were acquired as z-stacks with 1µM z-planes on Leica SP5 and SP8 confocal microscopes using a 40x 1.3 numerical aperture PL Apo Oil objective. All images were quantified using the PECAn image and data analysis pipeline (manuscript in preparation). Statistical analysis was performed using Rstudio and Graphpad Prism 8 software. Specific statistical tests and number of replicates performed for each experiment is provided in the statistical source data sheet. The following workflow was performed: if data met parametric assumptions (normality, homogeneity of variance) a t-test or paired t-test was used. If these criteria were not met, a Mann-Whitney U Test or Wilcoxon Signed-Rank test was used. A minimum of two independent biological replicates were performed of each experiment. Effect size metrics were performed only if a significant p-value was observed. Logistic regression was performed in PECAn. The dependent variable was the number of viable and non-viable *RpS3*^*+/-*^ cells in the loser clone border, as determined via a staining for cleaved DCP1. Predictor variables were determined by running the model with different, non-collinear variables (as determined by variance inflation factor below 5) and scoring by minimizing the Aikaike information criterion. An FDR correction was used to correct for multiple comparisons.

---

**Figure 1-figure supplement 1:**
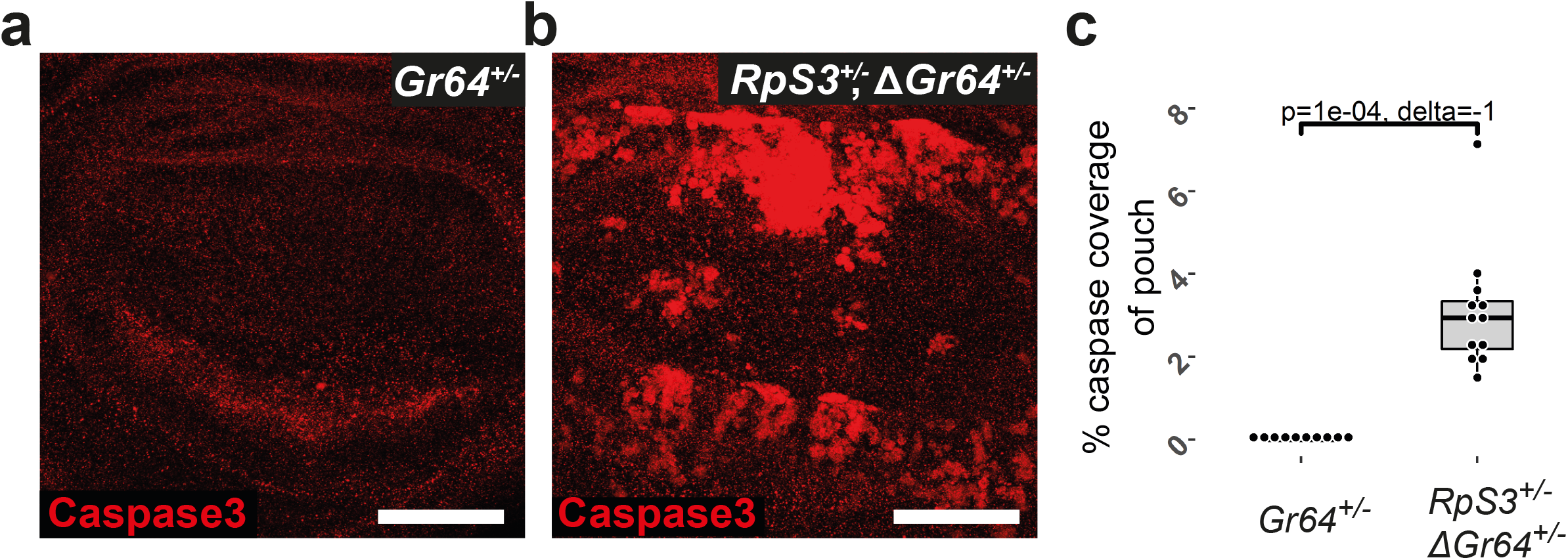
Effect of loss of *Gr64* on cell death in wildtype wing discs. Wing discs heterozygous for deficiency spanning the *Gr64* locus (ΔGr64) in a wildtype (**a**) or *RpS3*^*+/-*^ (**b**) background and assessed for cell death with a staining for cleaved-caspase3 (red), along with quantification in (**c**) (n _ΔGr64_=10, n_RpS3,ΔGr64_=11, two-sided Mann-Whitney U test with Cliff’s δ effect size metric).

**Figure 1-figure supplement 2:**
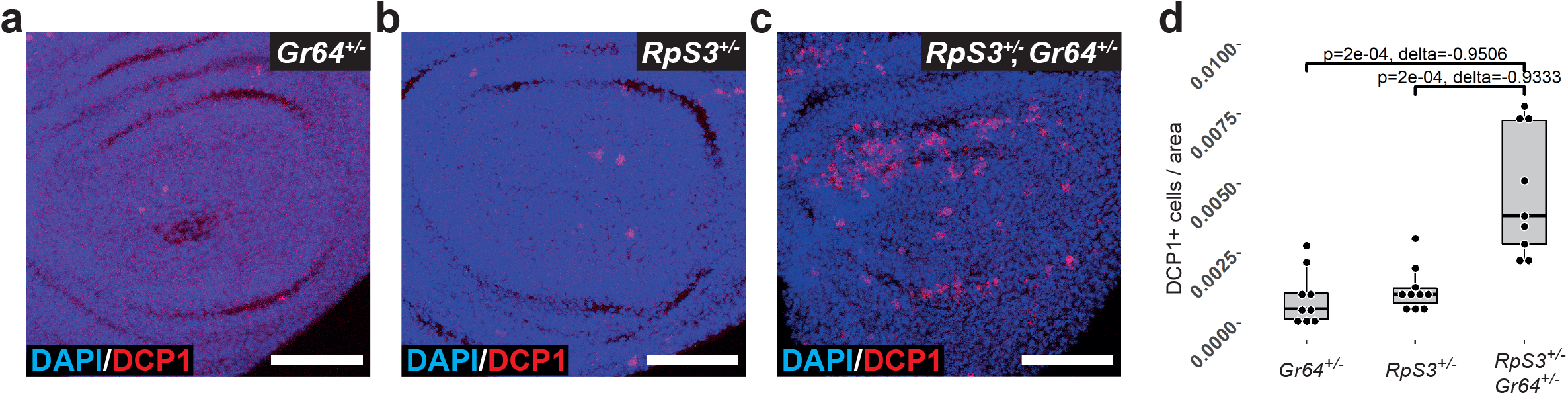
Confirmation of *RpS3*^*+/-*^, *Gr64*^*+/-*^ synthetic lethality using a precise *Gr64* cluster deletion. Wing discs heterozygous for a precise deletion of the *Gr64* cluster (**a**), heterozygous mutant for *RpS3*(**b**), or heterozygous mutant for both (**c**) assessed for cell death via a staining for cleaved-DCP1, along with quantification in (**d**) (n_Gr64_=9, n_RpS3_=10, n_RpS3,Gr64_=9, two-sided Mann-Whitney U test with Cliff’s δ effect size metric). Scale bars correspond to 50µM.

**Figure 1-figure supplement 3:**
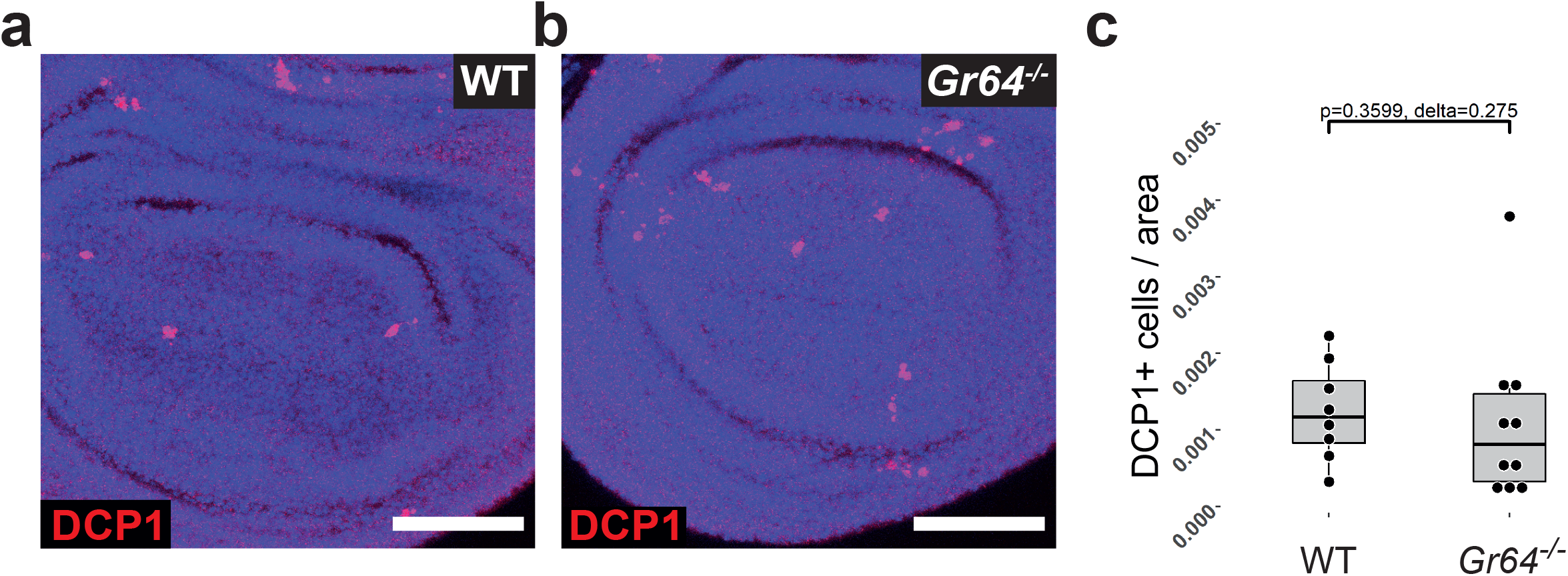
*Gr64* does not contribute to cell survival in the wildtype context. Wildtype (**a**) or *Gr64* homozygous null (**b**) wing discs assessed for cell death via a staining for cleaved-DCP1, along with quantification in (**c**) (n_wt_=8, n_Gr64_=10, two-sided Mann-Whitney U test with Cliff’s δ effect size metric). Scale bars correspond to 50µM.

**Figure 3-figure supplement 1:**
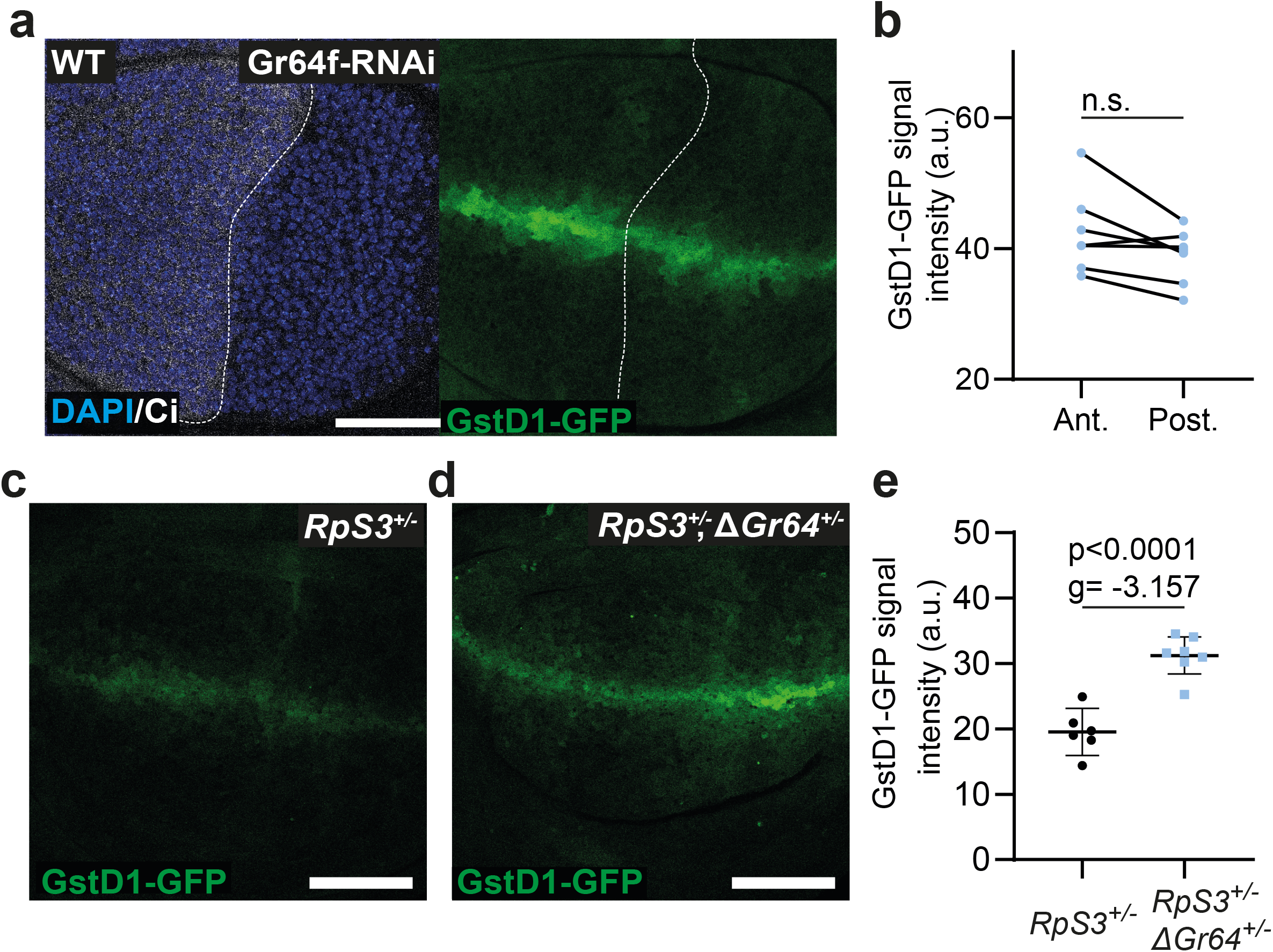
Effect of loss of Gr64 on oxidative stress activation in *RpS3*^*+/+*^ and *RpS3*^*+/-*^. (**a**-**b**) Wildtype wing discs expressing Gr64f-RNAi in the posterior compartment and assessed for GstD1-GFP reporter expression (green) along with quantification in (**b**) (n=7, two-sided t-test). (**c**-**e**) Wing discs heterozygous mutant for *RpS3* without (**c**) or with (**d**) a heterozygous deficiency in the *Gr64* locus (ΔGr64) and assessed for GstD1-GFP reporter expression (green) along with quantification in (**e**) (n_RpS3_=6, n_RpS3,ΔGr64_=7, two-sided student’s t-test with Hedges’ g effect size metric, the horizontal line indicates the mean and the whiskers reflect the 95% confidence interval).

**Figure 3-figure supplement 2:**
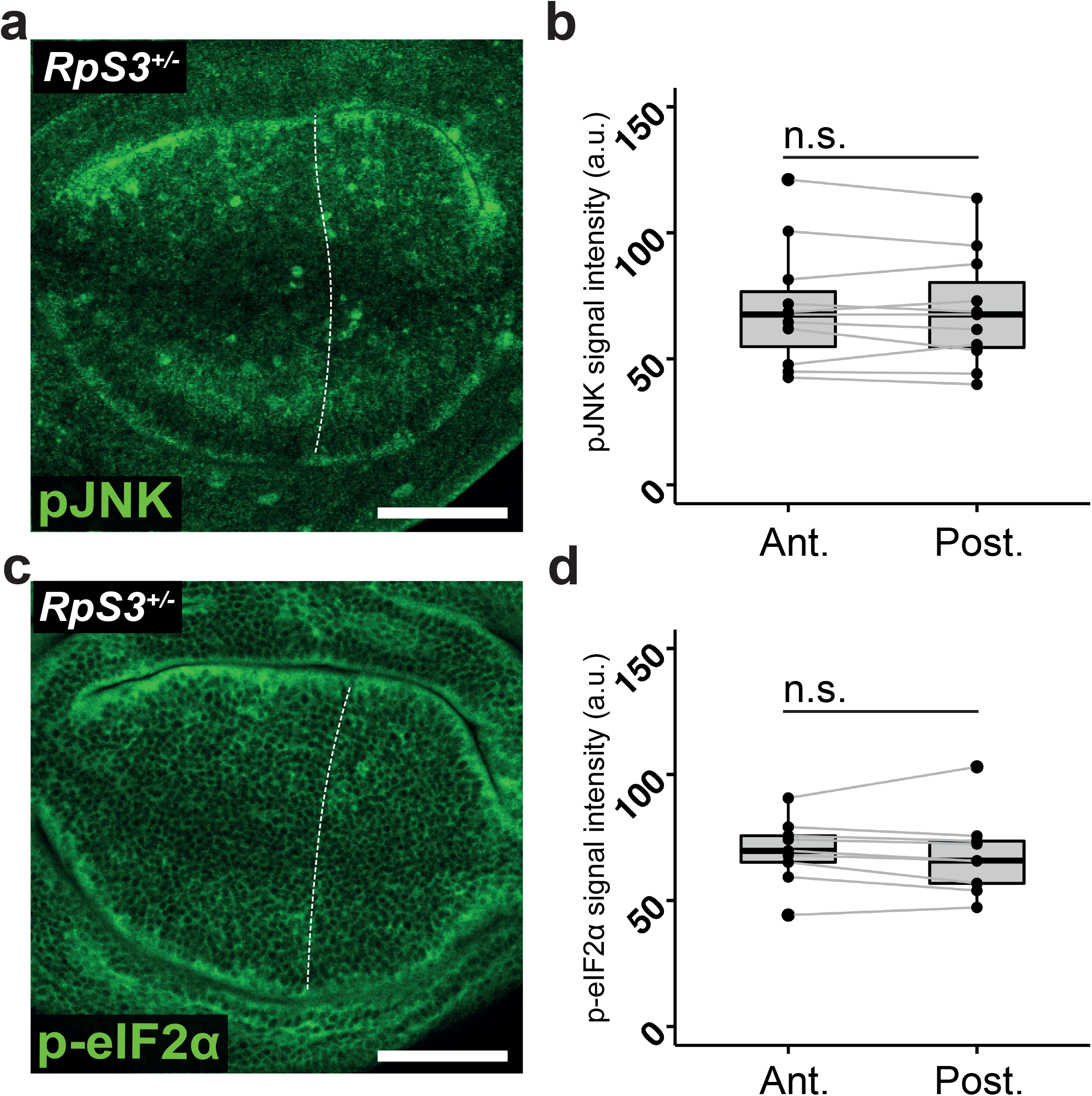
No RNAi negative controls for Figure 3. (**a**) *RpS3*^*+/-*^ wing discs carrying the *en-Gal4* driver but expressing no RNAi construct in the posterior compartment and stained for phosphorylated JNK (green) with quantification in (**b**) (n=11, two-sided paired t-test). (**c**) *RpS3*^*+/-*^ wing discs expressing the *en-Gal4* driver but no RNAi in the posterior compartment and stained for phosphorylated eIF2α (green) along with quantification in (**d**) (n=9, two-sided paired t-test).

## Works Cited

Albert, B., Kos-Braun, I. C., Henras, A. K., Dez, C., Rueda, M. P., Zhang, X., Gadal, O., Kos, M., & Shore, D. (2019). A ribosome assembly stress response regulates transcription to maintain proteome homeostasis. ELife, 8, e45002. https://doi.org/10.7554/eLife.45002

Armistead, J., & Triggs-Raine, B. (2014). Diverse diseases from a ubiquitous process: The ribosomopathy paradox. FEBS Letters, 588(9), 1491–1500. https://doi.org/10.1016/j.febslet.2014.03.024

Aspesi, A., & Ellis, S. R. (2019). Rare ribosomopathies: Insights into mechanisms of cancer. Nature Reviews Cancer, 19(4), 228–238. https://doi.org/10.1038/s41568-019-0105-0

Baillon, L., Germani, F., Rockel, C., Hilchenbach, J., & Basler, K. (2018). Xrp1 is a transcription factor required for cell competition-driven elimination of loser cells. Scientific Reports, 8(1), 17712. https://doi.org/10.1038/s41598-018-36277-4

Baumgartner, M. E., Dinan, M. P., Langton, P. F., Kucinski, I., & Piddini, E. (2021). Proteotoxic stress is a driver of the loser status and cell competition. Nature Cell Biology, 23(2), 136–146. https://doi.org/10.1038/s41556-020-00627-0

Blanco, J., Cooper, J. C., & Baker, N. E. (2020). Roles of C/EBP class bZip proteins in the growth and cell competition of Rp (‘Minute’) mutants in Drosophila. ELife, 9, e50535. https://doi.org/10.7554/eLife.50535

Bootman, M. D., Chehab, T., Bultynck, G., Parys, J. B., & Rietdorf, K. (2018). The regulation of autophagy by calcium signals: Do we have a consensus? Cell Calcium, 70, 32–46. https://doi.org/10.1016/j.ceca.2017.08.005

Bridges, C., & Morgan, T. (1923). The third-chromosome group of mutant characters of Drosophila Melanogaster. The Carnegie Institute of Washington.

Carreras-Sureda, A., Pihán, P., & Hetz, C. (2018). Calcium signaling at the endoplasmic reticulum: Fine-tuning stress responses. Cell Calcium, 70, 24–31. https://doi.org/10.1016/j.ceca.2017.08.004

Decuypere, J.-P., Bultynck, G., & Parys, J. B. (2011). A dual role for Ca2+ in autophagy regulation. Cell Calcium, 50(3), 242–250. https://doi.org/10.1016/j.ceca.2011.04.001

Ellis, S. R. (2014). Nucleolar stress in Diamond Blackfan anemia pathophysiology. Biochimica et Biophysica Acta (BBA) - Molecular Basis of Disease, 1842(6), 765–768. https://doi.org/10.1016/j.bbadis.2013.12.013

Ferrer, I., Garcia-Esparcia, P., Carmona, M., Carro, E., Aronica, E., Kovacs, G. G., Grison, A., & Gustincich, S. (2016). Olfactory Receptors in Non-Chemosensory Organs: The Nervous System in Health and Disease. Frontiers in Aging Neuroscience, 8. https://doi.org/10.3389/fnagi.2016.00163

Fujii, S., Yavuz, A., Slone, J., Jagge, C., Song, X., & Amrein, H. (2015). Drosophila Sugar Receptors in Sweet Taste Perception, Olfaction and Internal Nutrient Sensing. Current Biology : CB, 25(5), 621–627. https://doi.org/10.1016/j.cub.2014.12.058

Gaudel, F., Stephan, D., Landel, V., Sicard, G., Féron, F., & Guiraudie-Capraz, G. (2019). Expression of the Cerebral Olfactory Receptors Olfr110/111 and Olfr544 Is Altered During Aging and in Alzheimer’s Disease-Like Mice. Molecular Neurobiology, 56(3), 2057–2072. https://doi.org/10.1007/s12035-018-1196-4

Jiao, Y., Moon, S. J., Wang, X., Ren, Q., & Montell, C. (2008). Gr64f is Required in Combination with other Gustatory Receptors for Sugar Detection in Drosophila. Current Biology : CB, 18(22), 1797–1801. https://doi.org/10.1016/j.cub.2008.10.009

Katheder, N. S., Khezri, R., O’Farrell, F., Schultz, S. W., Jain, A., Rahman, M. M., Schink, K. O., Theodossiou, T. A., Johansen, T., Juhász, G., Bilder, D., Brech, A., Stenmark, H., & Rusten, T. E. (2017). Microenvironmental autophagy promotes tumour growth. Nature, 541(7637), 417–420. https://doi.org/10.1038/nature20815

Kim, H., Kim, H., Kwon, J. Y., Seo, J. T., Shin, D. M., & Moon, S. J. (2018). Drosophila Gr64e mediates fatty acid sensing via the phospholipase C pathway. PLOS Genetics. https://doi.org/10.1371/journal.pgen.1007229

Kucinski, I., Dinan, M., Kolahgar, G., & Piddini, E. (2017). Chronic activation of JNK JAK/STAT and oxidative stress signalling causes the loser cell status. Nature Communications, 8(1), 136. https://doi.org/10.1038/s41467-017-00145-y

Lee, C.-H., Kiparaki, M., Blanco, J., Folgado, V., Ji, Z., Kumar, A., Rimesso, G., & Baker, N. E. (2018). A Regulatory Response to Ribosomal Protein Mutations Controls Translation, Growth, and Cell Competition. Developmental Cell, 46(4), 456-469.e4. https://doi.org/10.1016/j.devcel.2018.07.003

Martín, F. A., Herrera, S. C., & Morata, G. (2009). Cell competition, growth and size control in the Drosophila wing imaginal disc. Development, 136(22), 3747–3756. https://doi.org/10.1242/dev.038406

Marygold, S. J., Roote, J., Reuter, G., Lambertsson, A., Ashburner, M., Millburn, G. H., Harrison, P. M., Yu, Z., Kenmochi, N., Kaufman, T. C., Leevers, S. J., & Cook, K. R. (2007). The ribosomal protein genes and Minute loci of Drosophila melanogaster. Genome Biology, 8(10), R216. https://doi.org/10.1186/gb-2007-8-10-r216

Meyer, S. N., Amoyel, M., Bergantiños, C., Cova, C. de la, Schertel, C., Basler, K., & Johnston, L. A. (2014). An ancient defense system eliminates unfit cells from developing tissues during cell competition. Science, 346(6214). https://doi.org/10.1126/science.1258236

Mills, E. W., & Green, R. (2017). Ribosomopathies: There’s strength in numbers. Science, 358(6363). https://doi.org/10.1126/science.aan2755

Miyamoto, T., & Amrein, H. (2014). Diverse roles for the Drosophila fructose sensor Gr43a. Fly, 8(1), 19–25. https://doi.org/10.4161/fly.27241

Miyamoto, T., Chen, Y., Slone, J., & Amrein, H. (2013). Identification of a Drosophila Glucose Receptor Using Ca2+ Imaging of Single Chemosensory Neurons. PLOS ONE, 8(2), e56304. https://doi.org/10.1371/journal.pone.0056304

Morata, G., & Ripoll, P. (1975). Minutes: Mutants of Drosophila autonomously affecting cell division rate. Developmental Biology, 42(2), 211–221. https://doi.org/10.1016/0012-1606(75)90330-9

Mukherjee, R., Das, A., Chakrabarti, S., & Chakrabarti, O. (2017). Calcium dependent regulation of protein ubiquitination – Interplay between E3 ligases and calcium binding proteins. Biochimica et Biophysica Acta (BBA) - Molecular Cell Research, 1864(7), 1227–1235. https://doi.org/10.1016/j.bbamcr.2017.03.001

Pakos-Zebrucka, K., Koryga, I., Mnich, K., Ljujic, M., Samali, A., & Gorman, A. M. (2016). The integrated stress response. EMBO Reports, 17(10), 1374–1395. https://doi.org/10.15252/embr.201642195

Park, J.-H., & Kwon, J. Y. (2011). Heterogeneous Expression of Drosophila Gustatory Receptors in Enteroendocrine Cells. PLOS ONE, 6(12), e29022. https://doi.org/10.1371/journal.pone.0029022

Recasens-Alvarez, C., Alexandre, C., Kirkpatrick, J., Nojima, H., Huels, D. J., Snijders, A. P., & Vincent, J.-P. (2021). Ribosomopathy-associated mutations cause proteotoxic stress that is alleviated by TOR inhibition. Nature Cell Biology, 23(2), 127–135. https://doi.org/10.1038/s41556-020-00626-1

Shirazi-Beechey, S. P., Daly, K., Al-Rammahi, M., Moran, A. W., & Bravo, D. (2014). Role of nutrient-sensing taste 1 receptor (T1R) family members in gastrointestinal chemosensing. The British Journal of Nutrition, 111 Suppl 1, S8–15. https://doi.org/10.1017/S0007114513002286

Slone, J., Daniels, J., & Amrein. (2007). Sugar Receptors in Drosophila. Current Biology, 17(20), 1809–1816. https://doi.org/10.1016/j.cub.2007.09.027

Sykiotis, G. P., & Bohmann, D. (2008). Keap1/Nrf2 Signaling Regulates Oxidative Stress Tolerance and Lifespan in Drosophila. Developmental Cell, 14(1), 76–85. https://doi.org/10.1016/j.devcel.2007.12.002

Tamori, Y., Bialucha, C. U., Tian, A. G., Kajita, M., Huang, Y.-C., Norman, M., Harrison, N., Poulton, J., Ivanovich, K., Disch, L., Liu, T., Deng, W. M., & Fujita, Y. (2010). Involvement of Lgl and Mahjong/VprBP in Cell Competition. PLOS Biology, 8(7), e1000422.

Tye, B. W., Commins, N., Ryazanova, L. V., Wühr, M., Springer, M., Pincus, D., & Churchman, L. S. (2019). Proteotoxicity from aberrant ribosome biogenesis compromises cell fitness. ELife, 8, e43002. https://doi.org/10.7554/eLife.43002

Yavuz, A., Jagge, C., Slone, J., & Amrein, H. (2015). A genetic tool kit for cellular and behavioral analyses of insect sugar receptors. Fly, 8(4), 189–196. https://doi.org/10.1080/19336934.2015.1050569

